# Advanced Deep Learning Enables Prediction of Allogeneic Stem Cell Mobilization Success

**DOI:** 10.1101/2025.09.17.676674

**Authors:** Asif Adil, Jingyu Xiang, Nicola Piccirillo, Hillary G. Harris, Simona Sica, John F. DiPersio, Stephanie N. Hurwitz

**Author notes:** Corresponding author: Stephanie N. Hurwitz, MD, PhD, Indiana University School of Medicine, Walther Hall, C321C, 980 West Walnut St, Indianapolis, IN 46202. **Competing interests** John F. DiPersio: Consulting/Advisory Committees Rivervest, Bioline, Incyte, NeoImmuneTech, Vertex, Bluebird Bio. Employment/salary: Washington University. Ownership Investment: Magenta, Wugen. All other authors declare no financial or non-financial competing interests.

## Abstract

Hematopoietic stem and progenitor cell (HSPC) transplantation offers a potentially curative therapy for aggressive hematologic malignancies and bone marrow failure syndromes. Successful transplantation depends on effective mobilization of donor CD34^+^ cells, yet some healthy donors fail to achieve adequate CD34^+^ yields despite standard granulocyte colony-stimulating factor (G-CSF)-based regimens. Early identification of such donors enables timely intervention, improving transplantation outcomes and reducing healthcare costs. We analyzed demographic and pre- and post-G-CSF laboratory data from 1,160 healthy donors from across multiple institutions and developed two complementary machine-learning frameworks to predict mobilization outcome. A transformer-based probabilistic model (TabPFN) trained on baseline complete blood counts (CBCs) rigorously discriminates poor from good mobilizers. Applying the same architecture to donor data after mobilization attains near-perfect discrimination. To unify the predictions across time points, we introduce an attention-aware neural network that ingests either baseline or post-mobilization data via a “lab-type” context flag, enabling accurate prediction of poor mobilizers both before and after GCSF mobilization. We further validated the framework on data from over 19,000 healthy donors compiled by the Center for International Blood and Marrow Transplant Research. These interpretable models enable early triage and “just-in-time” rescue interventions, providing a data driven foundation for personalized donor mobilization strategies.

## Introduction

Efficient mobilization of donor hematopoietic stem and progenitor cells (HSPCs) to peripheral blood (PB) is essential to the success of HSPC transplantation (HSPCT), a key intervention in the treatment of hematologic malignancies (1,2). Although granulocyte colony-stimulating factor (G-CSF) is widely used to induce peripheral mobilization of HSPCs, healthy allogeneic donors exhibit marked inter-individual variability in CD34^+^ cell yield (3–6). Inadequate mobilization can compromise stem cell collection, and there remains no reliable method to identify these individuals prior to apheresis collection (7). In routine practice, roughly 60-70% of donors achieve an optimal collection (5×10^6^ CD34^+^ cells/kg recipient weight) in one leukapheresis session, while the rest require multiple collections, and about 2–5% never attain the target dose (6,8). Suboptimal mobilization has serious clinical consequences, including delayed engraftment that leads to higher rates of graft failure, infection, bleeding, and relapse. Donors may also face complications of prolonged G-CSF treatment and multiple apheresis sessions (9). Overall, unpredictable donor mobilization remains a burden on healthcare cost and limits therapeutic success for HSPC transplant programs.

Known donor factors only partially account for this variability. Young age, male sex, and higher weight or body mass index (BMI) have been linked to better collection yields (6,10,11). However, a recent scoping review concluded that there is *“poor consensus”* on the best predictors of CD34^+^ yield (6), reflecting that most clinical studies report small or contradictory effects of individual covariates. In practice, donor registries and transplant centers largely rely on simple eligibility criteria (e.g. age limits) and basic laboratory tests (5).

In short, identifying poor mobilizers in advance remains challenging; most donors with borderline response are recognized only after mobilization has started. By that time, decisions about alternative strategies such as intensifying G-CSF dosing or adding plerixafor must be made reactively. Since remobilization or rescue interventions are expensive and time-consuming (4,12), there is a clear need for early prediction of mobilization outcomes to guide donor selection and apheresis planning.

Prior efforts to predict mobilization outcome have largely been confined to single-time-point analyses. Xiang et al previously combined baseline donor demographics with routine laboratory data to train machine-learning-based scoring models that identified suboptimal G-CSF mobilizers with approximately 83% accuracy (4). More recently, a multi-algorithm study used similar machine-learning models but reported only modest performance with best test-set accuracy <70 % and AUC <0.75 (12). A separate investigation categorized donors as “super” or “poor” mobilizers according to day 5 PB CD34^+^ cell concentrations and linked these categories to standard donor characteristics (13). Uniquely leveraging both baseline and post-mobilization data, Piccirillo et al used a multivariate logistic regression model to explain 26% of variability in mobilization, taking into account baseline platelet and white blood cell count (WBC) coupled with donor age and weight (5). Pre-G-CSF models, while useful for early risk stratification, may lack predictive resolution in borderline donors. Conversely, post-G-CSF models capture real-time biological response but are inherently reactive. A combined modeling strategy that leverages both pre- and post-G-CSF data offers the dual advantage of early prediction and refined risk assessment once G-CSF exposure has begun. This allows for a more nuanced understanding of donor mobilization dynamics and supports personalized decision-making throughout the mobilization timeline.

Building upon these, we utilize baseline and post-mobilization laboratory values to capture dynamic hematologic changes that occur over G-CSF dosing, identifying unique variables that define optimal donors at different timepoints. We demonstrate the performance of advanced deep learning models that can flexibly weigh routine laboratory data from both baseline and post-GCSF timepoints to accurately predict poor mobilizers, and validate an integrated model on data collected from over 19,000 donors in the Center for International Blood and Marrow Transplant Research (CIBMTR) database. Overall, these approaches identify donors at risk of suboptimal collection and support personalized mobilization strategies.

## Methods

### Study population and data acquisition

The study was approved by the Indiana University (IU) Institutional Review Board (IRB). Allogeneic HSPC donors from 2019-2024 at IU were identified (n=241). Donors were mobilized with G-CSF; a minor subset of patients additionally received a single dose of plerixafor. Donor demographic data and post-mobilization routine laboratory parameters, including complete blood count (CBC) and basic blood chemistries, were extracted from the electronic health record. All protected health information was de-identified before analysis in accordance with HIPAA regulations. During preprocessing, donors (n=38) with missing laboratory or demographic entries were removed, yielding the final curated IU dataset (n=203). To expand sample size and diversity, post-mobilization data from healthy donors (n=158) at the Università Cattolica del Sacro Cuore in Rome (CU) were combined with our patient population (5). These data were considered together as post-GCSF mobilized (total n=361).

To allow for additional comparison of the pre-mobilization timepoint, we compiled data from Washington University (WU) comprising 799 healthy donors who underwent baseline pre-G-CSF phlebotomy between 2010 and 2022 (4), forming the study’s pre-G-CSF donor data.

To further test the generalizability of our attention aware framework, we utilized a publicly available donor cohort (n=22,348) from the CIBMTR, including both pre- and post-GSF laboratory values with donor demographics (14). The dataset is part of a broader registry maintained by the CIBMTR, a collaborative initiative between the Medical College of Wisconsin and the National Marrow Donor Program that collects comprehensive clinical data from more than 300 transplant centers globally. Donors with incomplete laboratory records or missing BMI values were excluded. Post-mobilization laboratory measurements were restricted to those obtained on the first day of collection, resulting in a final analytic cohort of 19,207 donors.

### Mobilization threshold definition

To establish a binary definition of mobilization quality independent of recipient variables, donor PB CD34^+^ cell/μL measured on the day of collection was compared to total CD34^+^ cells/recipient kilogram (kg) weight collected. This analysis was performed using the IU donor subset mobilized with GCSF alone (n = 170). A strong positive correlation between these values was observed, supporting the use of CD34^+^ cell/μL as a donor-dependent outcome metric, as previously described (4,15). Based on total CD34^+^ cells/recipient kilogram weight, donors were categorized into “good” (≥5 ×10^6^), and “poor” (<5 ×10^6^). Median yields were calculated for each category of donors to establish the cutoff of ≥40 CD34^+^ cells/μL to define “good mobilizers,” a threshold that is consistent with that used previously (4).

## Results

### Defining a mobilization threshold and constructing the integrated donor cohort

To develop and evaluate predictive models for stem cell mobilization outcomes, we implemented a structured and reproducible analytical pipeline (**Fig. 1**).

**Figure 1.**
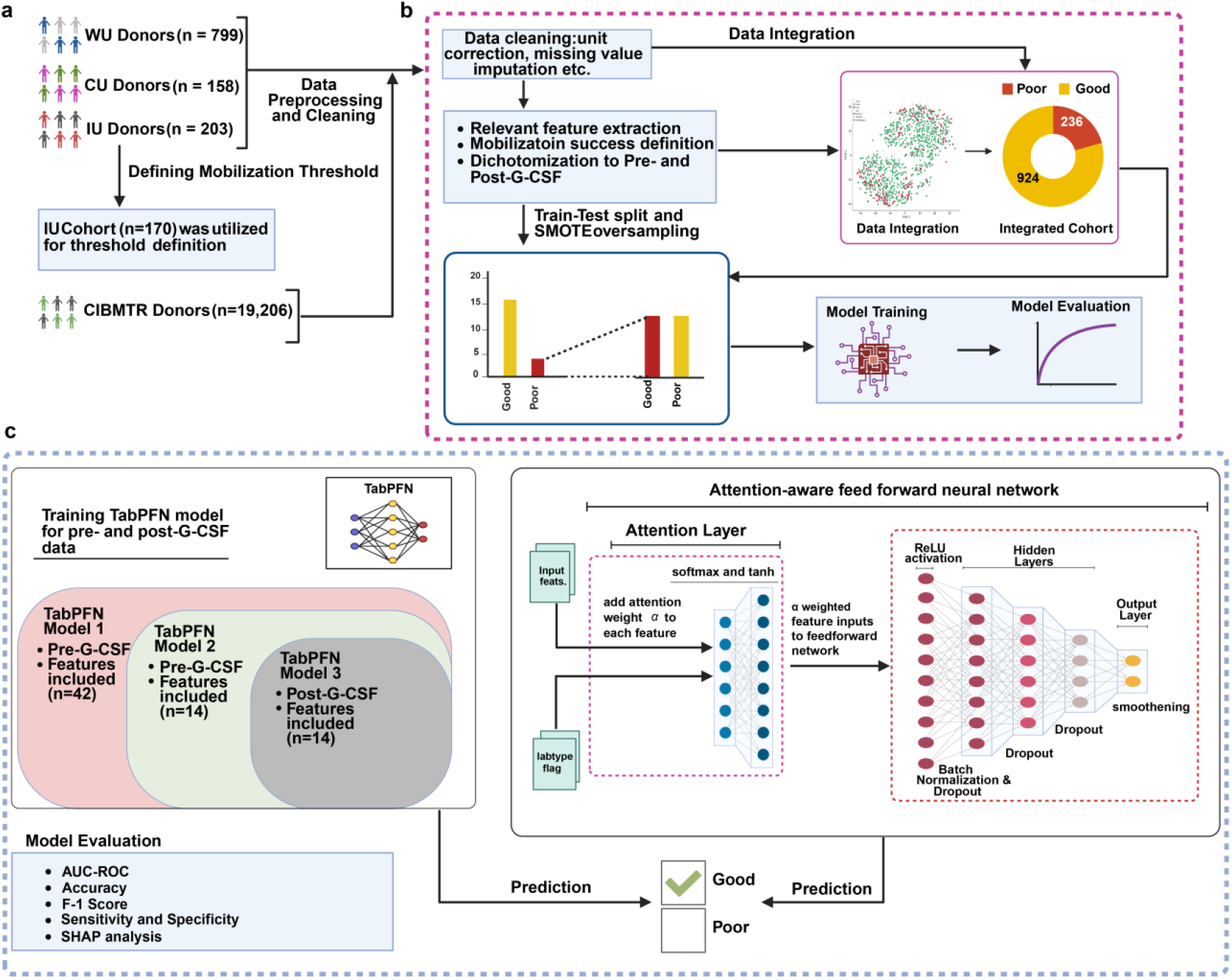
Schematic overview of the analytical pipeline. **a)** Data acquisition and preprocessing. Peripheral blood counts and demographics were collated from three independent centers (Washington University, WU; Catholic University, CU; Indiana University, IU and CIBMTR, Center for International Blood and Marrow Transplant Research). An empirical mobilization threshold was first defined in an IU subset (n = 170). **b**) All cohorts then underwent unit harmonization, missing-value imputation, and SMOTE-based oversampling to correct class imbalance before an 80:20 stratified train/test split. After batch correction, the three datasets were merged into an integrated cohort (good mobilizers = 924, poor mobilizers = 236) for exploratory analyses. **c)** Model development and evaluation. Left: Three TabPFN models were trained: (1) full pre-G-CSF feature set (42 variables); (2) reduced pre-G-CSF panel (14 variables); and (3) post-G-CSF panel (14 variables). Right: An attention-aware feed-forward neural network leveraging the “lab-type” domain indicator was trained in parallel. Each model outputs a binary prediction (good vs poor) that is assessed by precision–recall curves, conventional performance metrics, and XAI-based feature attribution.

Data from G-CSF-mobilized IU donors were used to quantitatively define a “good” mobilizer based on a prior strategy (4). Overall, PB CD34^+^ cell/μL concentrations on the day of collection showed a positive correlation with the total CD34^+^ cell/recipient kg yield from apheresis (r = 0.47, p < 0.0001; **Fig. 2a**). Median CD34^+^ cell yields (x 10^6^/recipient kg) were calculated for donors below or above the standard threshold of 5 x 10^6^ CD34^+^ cells/kg to yield a cutoff of ≥ 40 CD34^+^/μL for defining “good mobilizers” (**Fig. 2b**).

**Figure 2.**
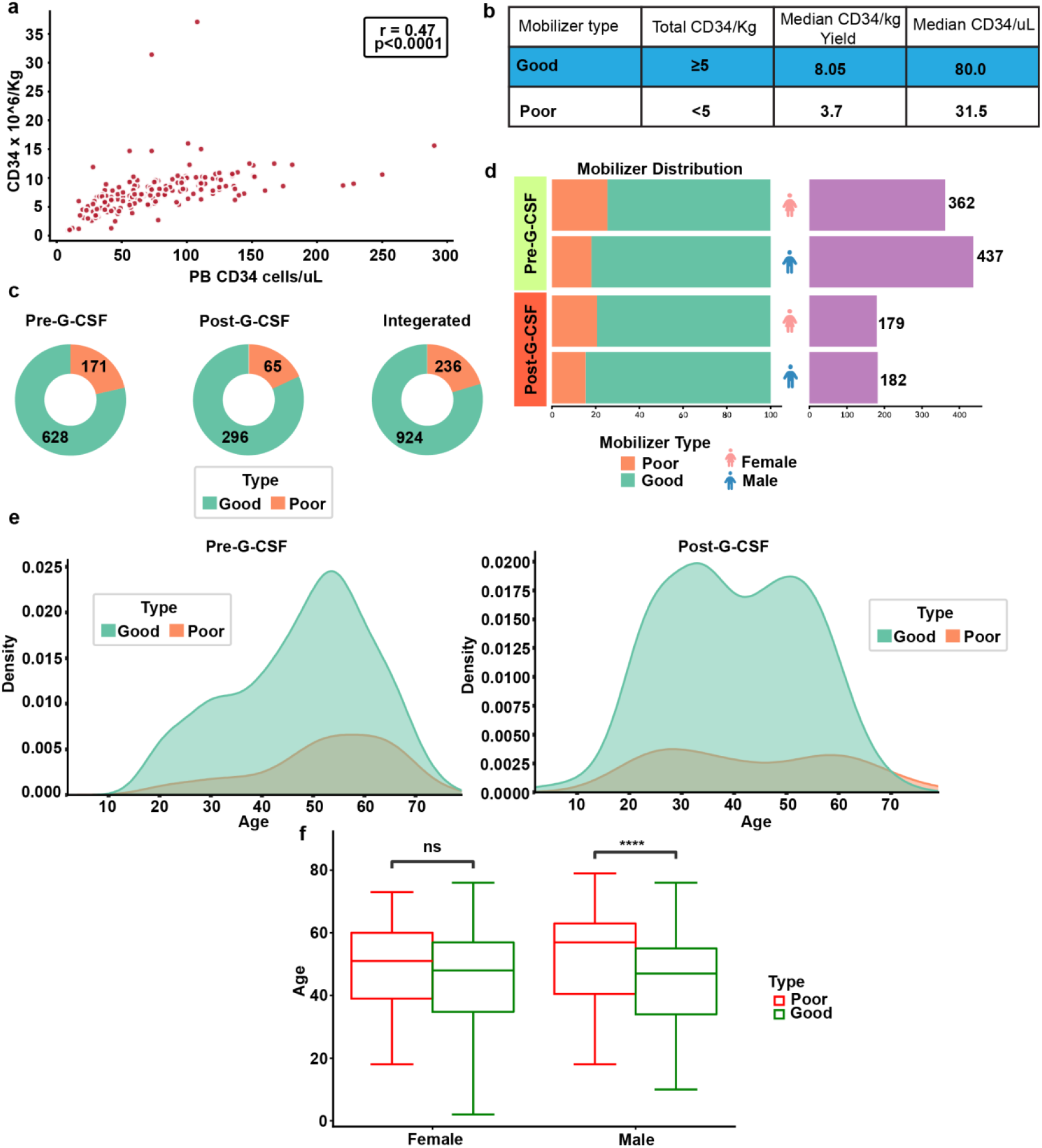
Characterization of donor mobilization efficiency across G-CSF timepoints and associated demographic patterns. **a)** Correlation of IU GCSF-mobilized donor peripheral blood CD34^+^ counts (day of collection) with total CD34^+^ cell/recipient kg yield (r = 0.47, p < 0.0001). **b)** Table showing the median values of Day 1 PB CD34^+^ count and yield across mobilizer categories. **c)** Frequency of good and poor mobilizers in the pre- and post-G-CSF cohorts, based on a 40 CD34^+^ cell/μL threshold. **d)** Distribution of good and poor mobilizers across cohorts and by sex. **e)** Kernel-density plots of donor age across mobilizer type in the pre- and post-G-CSF cohorts. **f)** Box-and-whisker plots comparing age and sex as mobilization factors. ****, *p* < 0.0001; ns, not significant.

Applying this threshold to the combined multi-institutional dataset, the majority of donors (∼80% overall) met the criteria for good mobilization (**Fig. 2c**). Frequencies of suboptimal/poor mobilization align with prior observations that roughly 10–20% of healthy donors can exhibit inadequate CD34^+^ mobilization after G-CSF stimulation (16,17).

Similar to prior reports, female donors were slightly overrepresented among the poor mobilizers compared to male donors (**Fig. 2d)** (16). Overall older donors exhibited poor mobilization (**Fig. 2e**). However, similar to prior reports, donor sex by itself was not an independent predictor of mobilization success in our study (5,18). When stratified by both sex and age, we note that male donors who were poor mobilizers were significantly older than male donors who mobilized successfully (p < 0.0001) (**Fig. 2f**). By contrast, among female donors, there was no significant age difference between poor and good mobilizers. These data suggest that advanced age may be a key contributor to mobilization inefficiency in the male population.

### Predicting mobilization outcome using TabPFN and pre-G-CSF donor features

Building on a prior prediction model developed by Xiang et al (4), we applied a transformer-based probabilistic model (TabPFN) (19) to demographic and baseline laboratory data from 799 healthy donors prior to G-CSF administration. For our model, input features included standard CBC indices, basic blood chemistry values, and donor demographics such as age, sex, and BMI.

This model achieved high predictive performance, with an accuracy of 89%, Mathews Correlation Coefficient (MCC) of 0.79 and AUC of 0.96 on the test set (**Fig. 3a-b, Supplementary Methods**).

**Figure 3.**
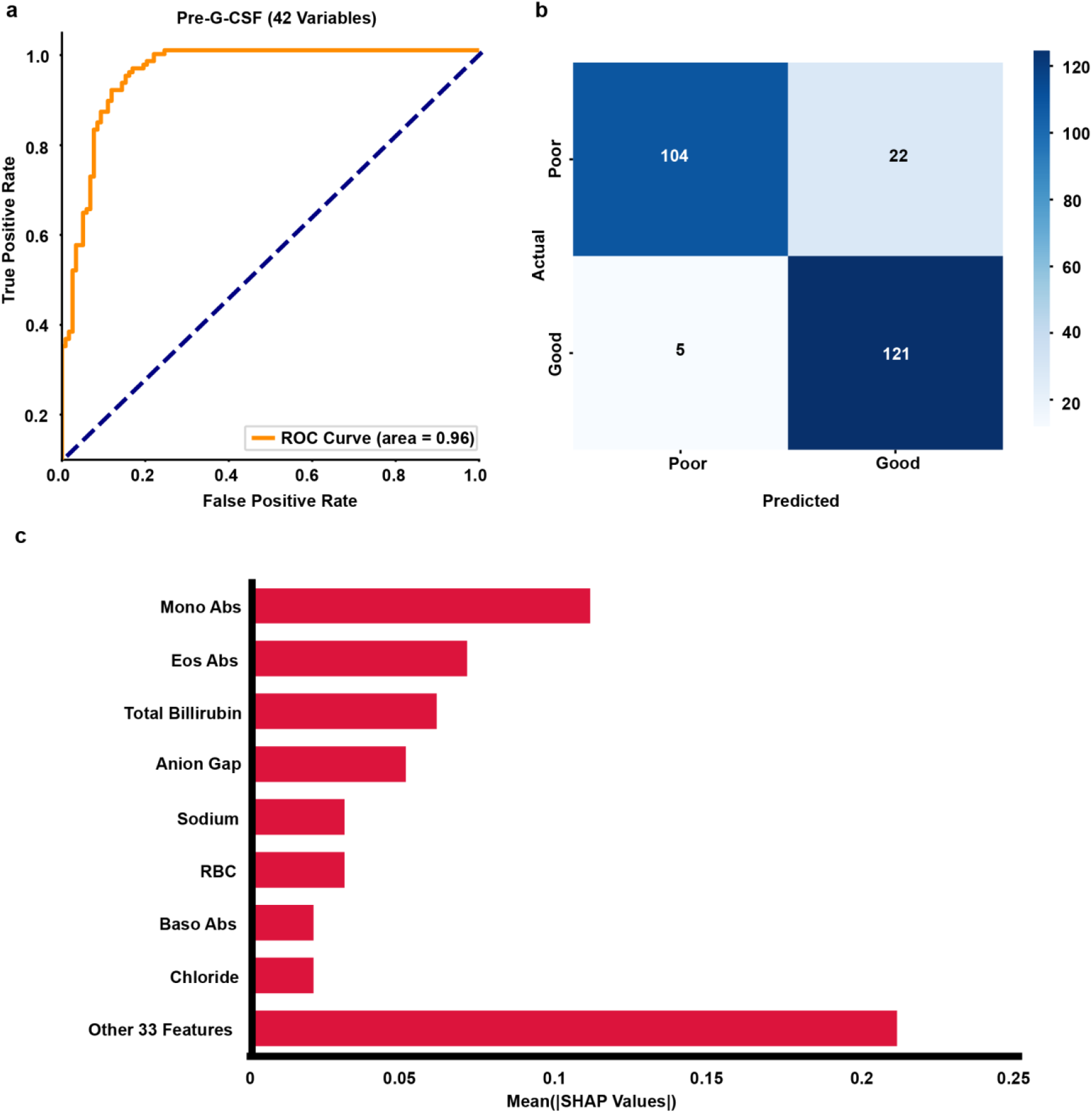
TabPFN performance on pre-G-CSF donor data. **a)** ROC curve for the held-out test set showing an area under the curve (AUC) of 0.96. The dashed line denotes random performance. **b)** Confusion matrix depicting that the model correctly classified 219 (overall accuracy = 87%), with 94 true negatives, 125 true positives, 29 false positives and 4 false negatives, reflecting sensitivity of 97%. Cell shading is proportional to count. **c)** SHAP summary plot depicting the contribution of each input variable to the model’s output, ordered by mean absolute SHAP value (model impact).

SHapley Additive Explanations (SHAP) interpretability analysis indicated that predictive importance was concentrated in age, BMI, and conventional hematological indices: absolute monocyte, eosinophil, and basophil counts; and erythrocyte and platelet parameters (**Fig. 3c**). In contrast, biochemical and coagulation variables contributed only a modest, diffuse signal. These data align with clinical observations that higher pre-apheresis monocyte counts and platelet indices are independently associated with successful CD34^+^ collections, whereas advancing age, lower BMI, and higher MCV correlate with inferior yields (9,20–24).

### Performance of TabPFN using baseline CBC and demographic features alone

To determine whether routinely collected demographic and pre-mobilization CBC data are sufficient for accurate risk stratification, we retrained TabPFN using only 14 baseline variables: CBC indices, donor age, and sex. Despite the substantial reduction in feature breadth, the model retained excellent discriminative ability, achieving an accuracy of 91%, MCC = 0.83 and AUC of 0.96 on the held-out test set (**Fig. 4a-b**). The AUC was similar to the full 42-feature panel, indicating that the primary signals for distinguishing mobilizer status come from baseline hematology and demographics.

**Figure 4.**
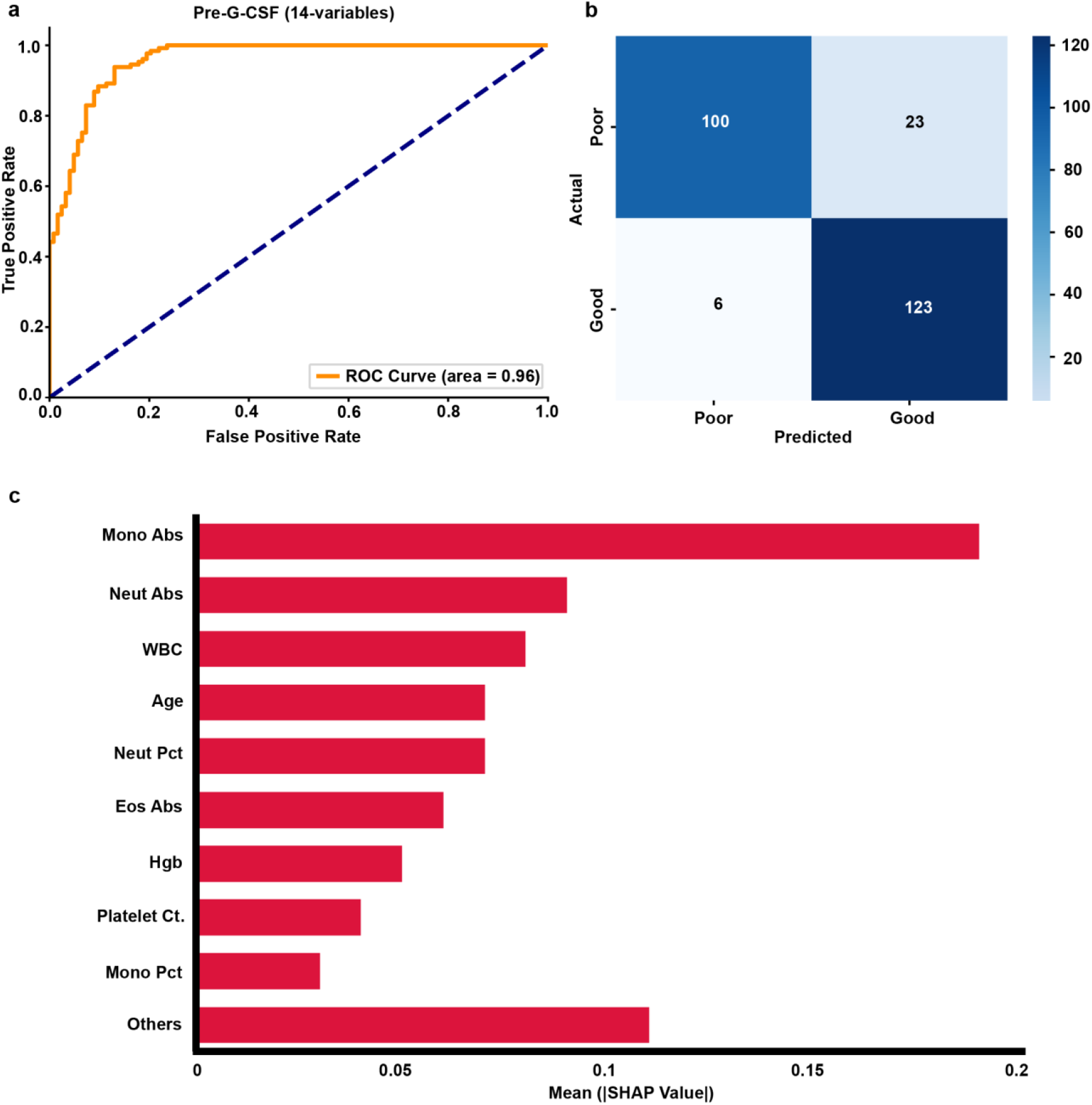
TabPFN performance using limited features of pre-GCSF donor data. **a)** ROC curve for the independent test set showing an area under the curve (AUC) of 0.96. The dashed line denotes random performance. **b)** Confusion matrix summarizing predictions of 252 donors with overall accuracy of 89%, sensitivity of 95% (123 / 129 positives), and specificity of 81% (100 / 123 negatives); 23 false positives and 6 false negatives account for the residual error. Cell shading is proportional to count. **c)** SHAP summary plot depicting the contribution of each input variable to the model’s output, ordered by mean absolute SHAP value (model impact).

Interpretability analysis corroborated known clinical correlates of mobilization success (**Fig 4c**). Consistent with earlier observations, monocytes, neutrophils, WBC, and age emerged as the most influential variables. These results align with previous reports that advanced age and lower baseline leukocyte parameters predict reduced CD34^+^ yields after G-CSF stimulation (16,17), strengthening confidence that the observed relationships are biologically meaningful.

### Post-G-CSF prediction of mobilization outcome using TabPFN

Using the same 14 input features (**Supplementary Table 1)** TabPFN demonstrated excellent performance on donor data in the post-G-CSF mobilization cohort. It achieved near-perfect discrimination, with an accuracy of 94%, MCC=0.90 and ROC AUC of 0.99, slightly outperforming the pre-G-CSF model, and showing 93% sensitivity and 96% specificity (**Fig. 4a-b**).

SHAP analysis of the post-G-CSF model revealed that neutrophil percentage was the single most predictive feature, followed by hematocrit, lymphocyte count, hemoglobin, total WBC, and donor age (**Fig. 5c**). Notably, monocyte count was not considered a key predictive feature in the post-mobilization donor data. Overall, feature importance ranking mostly mirrored the pre-G-CSF model’s findings, indicating that G-CSF administration amplified, but did not substantially alter the relative importance of key hematologic and demographic predictors. This unbiased, direct model-to-model comparison of feature importance suggests that donor demographic and CBC indices alone may be used to predict donor mobilization success, irrespective of the evaluation timepoint.

**Figure 5.**
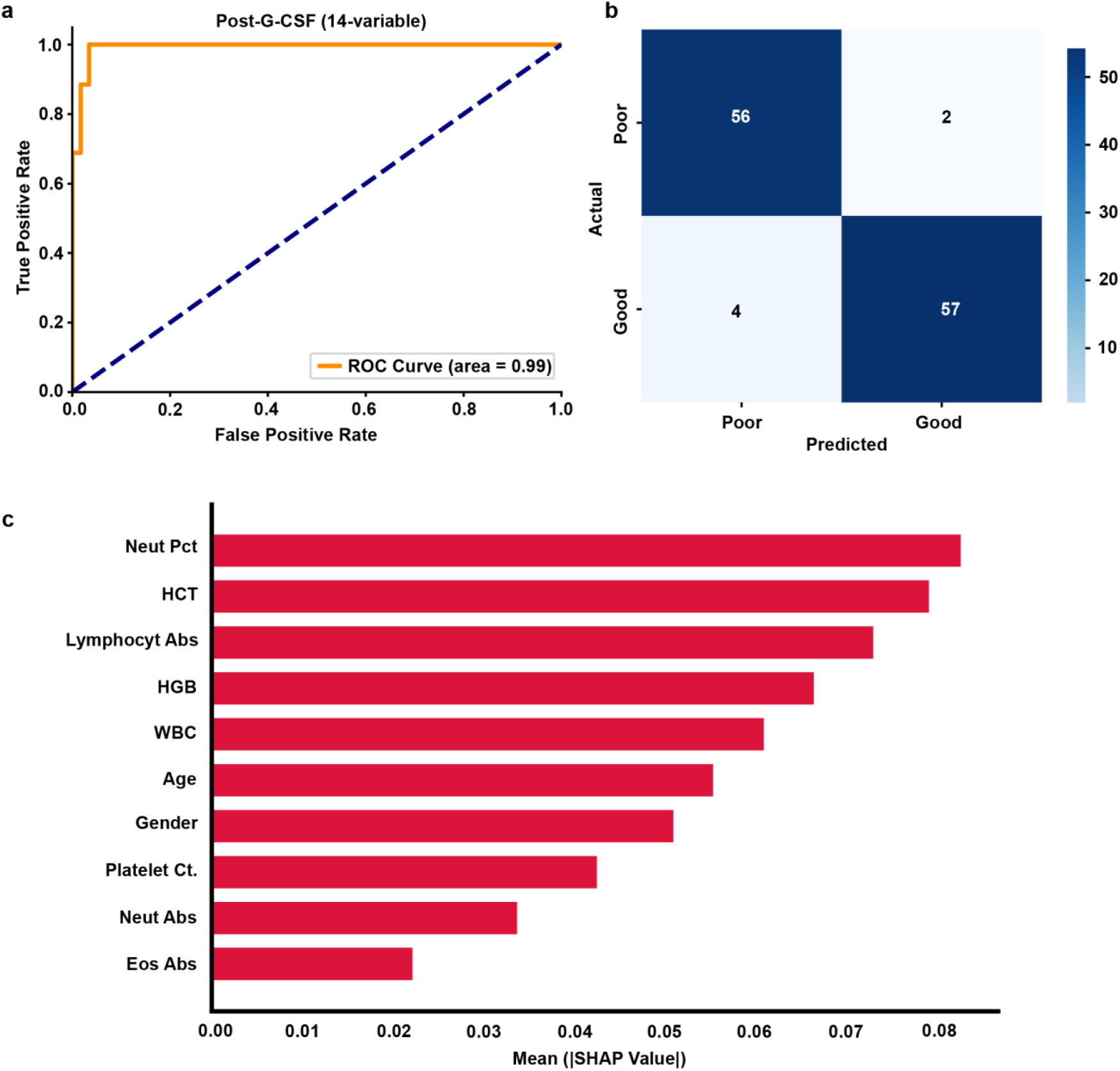
TabPFN performance on post-G-CSF donor data. **a)** ROC curve for the held-out test set showing an area under the curve (AUC) of 0.99. **b)** Confusion matrix of 119 donors showing an overall accuracy of 95%, sensitivity of 93 % (57/61) and specificity of 97% (56/58). Cell shading corresponds to count. c) SHAP summary plot depicting the contribution of each input variable to the model’s output, ordered by mean absolute SHAP value (model impact).

### Predicting mobilization outcome using an attention-aware deep learning model across temporal lab contexts

To leverage the full cohort of 1,160 donor samples spanning both pre- and post-G-CSF CBCs, we trained an attention-aware deep learning model that incorporates data from both timepoints (**Fig. 6a, Supplementary Methods**). The input feature vector included each donor’s CBC measurements and demographic variables (age, sex), with a binary lab-type flag (pre-G-CSF vs post-G-CSF) added to indicate the sample’s context. This unified model was designed to predict the mobilization outcome regardless of whether a donor’s CBC values were measured before or after G-CSF administration. By integrating the pre- and post-G-CSF data with an explicit context indicator, the model could learn shared predictive patterns while still capturing the distinct hematologic profiles characteristic of each condition.

**Figure 6.**
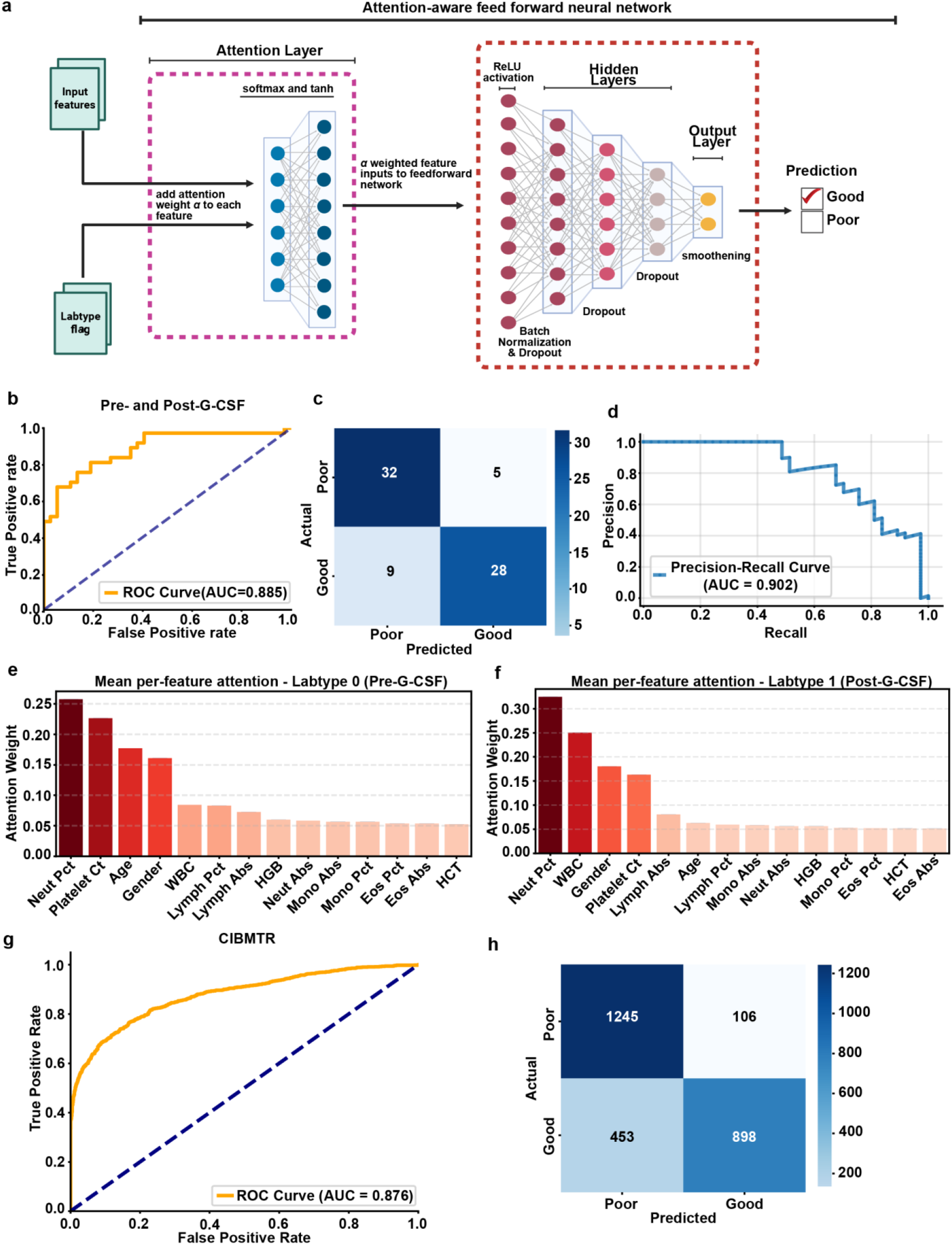
Attention-aware deep learning unified model performance on donor data across timepoints. **a)** Attention-aware feed-forward neural network for donor mobilization prediction. Clinical and complete blood count variables, together with a binary pre-/post-G-CSF flag (Labtype), enter an attention gate that assigns a normalized weight (α) to each feature via a soft-max + tanh mechanism. The re-weighted feature vector is passed to a multi-layer perceptron that incorporates ReLU activation, batch normalization, and dropout regularization. A final sigmoid node outputs the probability of successful (“good”) versus inadequate (“poor”) CD34^+^ cell mobilization, while the learned α weights are retained for downstream interpretation of driver features. **b)** ROC curve for the held-out test set depicts an AUC of 0.88. **c**) Confusion matrix for 74 donor test set showing 59/74 correct (accuracy = 81%), with a sensitivity of 84% (31/37) and specificity of 76% (28/37). **d)** Precision-recall curve illustrating model performance across classification thresholds, maintaining a precision >0.80 across most recall values. Mean self-attention weights for samples collected **e)** before GCSF (labtype 0) or **f)** after GCSF administration (labtype 1). **g)** ROC curve for held-out test depicts an AUC of 0.87 for the model trained on CIBMTR donor data of over 19,000 healthy donors. **h)** Confusion matrix for the model evaluated on the CIBMTR dataset (held-out test set, n = 2702) showing an accuracy of 79%, sensitivity of 66% (898/1,351) and specificity of 92% (1,245/1,351).

On evaluation, the model achieved an accuracy of 81%, MCC = 0.64, with a precision of 0.85 and a recall of 0.76, yielding a balanced F1 score of 0.80 (**Fig. 6b-c**). The AUC was 0.89, reflecting strong overall discriminative ability. Across different classification thresholds, the model tended to maintain a precision above 0.80 for different recall values (**Fig. 6d**).

We also analyzed the attention-based feature weights to understand which inputs the model found most predictive in each context. This revealed distinct importance patterns for pre- vs post-G-CSF samples. For pre-G-CSF cases (**Fig. 6e**), the model placed greatest weight on age and neutrophil percentage, followed by sex, lymphocyte percentage, and total WBC. For post-G-CSF cases (**Fig. 6f**), the highest weighted features were total WBC count and neutrophil percentage, with sex and age contributing next prominently.

To further evaluate the generalizability and clinical relevance of the framework, we applied it to a large CIBMTR dataset of healthy donors with pre- and post-G-CSF laboratory values. Despite using only eight reported features (hemoglobin, platelet count, WBC, absolute neutrophils, MNC, age group, BMI group, and weight), the model achieved an AUC of 0.88 (**Fig. 6g)** and an accuracy of 79% (**Fig. 6h)**. Attention weights of variables remained consistent with prior models (**Supplementary Fig. 1a-b**). Finally, attention entropy analysis and Jensen–Shannon divergence between attention distributions confirmed meaningful adaptation to the pre- or post-GCSF context while maintaining shared predictive structure (**Supplementary Fig. 2a-b, Supplementary Results**).

## Discussion

This study evaluates the performance of transformer-based and attention-aware deep learning models for HSPC mobilization outcomes using pre- and post-mobilization laboratory data. We found that (i) TabPFN achieves excellent discrimination with baseline CBC and demographic features (AUC = 0.96), (ii) performance was retained with only 14 core variables (AUC = 0.96), and (iii) post-mobilization data further improves accuracy (AUC = 0.99). Finally, our unified attention-aware model, integrating both timepoints via a lab-type flag and feature-level attention, produces robust discrimination (AUC = 0.89; F1 = 0.80) while offering interpretable context-specific feature weighting. Notably, the attention-aware model yielded a slightly lower AUC, reflecting the challenge of integrating heterogeneous pre- and post-mobilization features. In other words, once G-CSF has begun, the peripheral blood counts clearly indicate mobilization potential, whereas a model that tries to fuse pre- and post-data must weight signals across time and requires reconciling conflicting signals. Nevertheless, all models showed good discrimination, implying that routine demographic and CBC features alone may predict mobilization success with high accuracy.

The outputs align with current biological understanding. Older age and lower baseline platelet or leukocyte reserves are recognized impediments to CD34^+^ mobilization (18), whereas higher hematocrit, neutrophil percentage, and total WBC correlate with greater stem-cell yield (5,25). Our analyses revealed that pre-G-CSF importance is dominated by age and baseline innate immune cell counts, whereas post-G-CSF importance shifts toward the magnitude of the leukocyte surge (minimizing age as a variable). This supports that baseline hematologic reserve and dynamic marrow responsiveness jointly determine mobilization efficiency. The observation that routine chemistries are dispensable for predictive models echoes a 2024 scoping review, which found no serum-chemistry or coagulation marker with reproducible association after adjustment for age and blood counts (6), and a multivariable analysis of 84 sibling donors in which only hematologic parameters remained significant (26).

Furthermore, the attention-aware model provides an integrated solution for mixed time-points by re-weighting features according to the lab context flag. This flexibility is advantageous in real-world settings where centers collect laboratory data at different stages of donor evaluation and mobilization and wish to apply a single decision tool across time. More dynamic laboratory testing over various timepoints during mobilization is also certainly possible in future clinical trials, where this model could be applied. To test this generalizability, we applied our attention-aware framework to a large, publicly available dataset of over 19,000 healthy donors compiled by the CIBMTR (14). Despite differences in feature definitions and population composition, the model achieved an accuracy of 79%, demonstrating its applicability across diverse real-world donor populations. Of note, this dataset included only eight routine features, which likely limited the model’s discriminative capacity relative to our institutional dataset. Importantly, attention weight patterns remained consistent with biological expectations and previous findings, reinforcing the model’s capacity to prioritize clinically meaningful features across mobilization phases.

Clinically, a high-fidelity baseline donor predictor enables early donor triage, facilitating the selection of alternative donors or modification of mobilization regimen, such as intensified G-CSF dosing or addition of plerixafor. Piccirillo et al. proposed a simple age and WBC score that stratifies donors into 4% versus 46% risk of sub-optimal mobilization; our transformer model automates and refines this concept at scale (5). Conversely, the post-G-CSF model supports “just-in-time” rescue. For example, in a prospective phase-II study, a single plerixafor dose administered to donors with <2 × 106 CD34^+^ cells/kg after the first apheresis raised median peripheral CD34^+^ counts three-fold and enabled 57% of poor mobilizers to reach target yield the next day (27); similar interventions have demonstrated significant healthcare cost reduction for patients and hospitals (28). It is also worth noting recent evidence from our lab and others supporting better overall survival with higher CD34^+^ doses (>7 x 10^6^/kg) in certain patient populations, such as allogeneic matched sibling donors (29). These studies indicate that the frequency of suboptimal donors may be higher than estimated in this study, and add to the growing importance of identifying donors who are unlikely to meet these optimal dose collections.

Growing interest in the development of short-acting mobilizing agents also aims to provide alternative or rescue approaches to improve donor cell collection at the first apheresis attempt. In preclinical and clinical studies, motixafortide (BL-8040, BKT140), a selective CXCR4 antagonist rapidly increased PB CD34^+^ cell counts, peaking at 2-4 hours after the first dose (30,31) (32). Very late antigen 4 (VLA4) inhibitors (BIO5192, BOP) and a CXCR2 agonist (MGTA-145/GROβT) have also demonstrated efficacy as rapid mobilizing strategies (33–35). In the future, it is likely that incorporation of donor mobilization prediction strategies coupled with rapid rescue mobilization agents will reduce failed collections, shorten mobilization courses, and curb unnecessary apheresis sessions. By aligning high-accuracy model outputs with established biology, demonstrating tangible clinical use-cases, we offer a foundation for prospective validation and eventual integration of data-driven decision support into HSPC donor management.

## Acknowledgements

The authors thank the IU Division of Transfusion Medicine for assistance in compiling institutional donor data.

## Author contributions

Conceptualization: A.A. and S.N.H.; Data Curation: A.A., J.X., N.P., H.G.H., S.S., J.F.D., S.N.H.; Investigation and Formal analysis: A.A., Methodology: A.A.; Supervision: S.N.H.; Writing-original draft: A.A. and S.N.H.; Writing-review & editing: A.A., J.X., N.P., H.G.H., S.S., J.F.D., S.N.H.

## Data availability

Data supporting the findings of this study can be obtained from the corresponding author upon reasonable request.

## Code availability

All codes and trained models are available at: https://www.github.com/asif7adil/deepmop

