## Supplemental material for "Advanced Deep Learning Enables Prediction of Allogeneic Stem Cell Mobilization Success"

[#] Corresponding author:

Stephanie N. Hurwitz, MD, PhD

**Supplementary Methods:**
**Data integration for the attention aware deep learning model**

To generate the unified dataset required for training our self attention–aware deep learning model, donor records from IU, WU, and CU were merged, including 14 variables common to all cohorts: 12 CBC indices, donor age, and biologic sex. Continuous variables were scaled to the 0–1 interval using min–max normalization.

We performed exploratory analyses on the integrated dataset to characterize feature distributions and relationships. In addition, summary statistics were generated for donor subgroups, and two-tailed Student's t-tests were used to compare key features between good vs. poor mobilizer groups. All the results were considered significant if p-value was <0.05.

**Feature engineering**

To enhance the predictive performance of models trained on datasets with a limited number of input features, we employed several feature engineering strategies, including one-hot encoding (1), feature derivation, and imputation techniques. One-hot encoding was applied to categorical variables such as "Age-group" and "BMI-group" in the CIBMTR donor dataset (2) , allowing the model to interpret group-based differences without imposing ordinal assumptions. Additionally, derived features were constructed to enrich the biological relevance of the input space and to address missing values using clinically valid relationships. For example, neutrophil percentage was calculated from total WBC and absolute neutrophil count when not explicitly available. These computed values allowed for consistent representation across donors and preserved important immunological signals relevant to mobilization biology. All numerical features, including derived variables, were standardized prior to model training to ensure numerical stability and prevent feature scale bias.

**Handling class imbalance**

Because the datasets exhibited class imbalance (disproportionate numbers of good vs. poor mobilizers), we applied the Synthetic Minority Oversampling Technique (SMOTE) (3) to the training data to balance class frequencies. SMOTE generates synthetic minority-class examples by interpolating between existing minority samples. This oversampling approach was restricted to the model training phase to avoid information leakage. By achieving an approximately 1:1 class ratio in the training set, we aimed to prevent biased model learning and improve generalization to both mobilization outcomes.

**Attention aware deep-learning architecture**

A self-attention–aware feed-forward network was implemented in TensorFlow/Keras to classify donors as good or poor mobilizers. The model accepts two inputs: **feature vector (***feat_in*), **lab-type flag** (*lab_in*). To allow the network to weight predictors differently according to sampling context, we introduced a *FeatureAttention* layer which concatenates the feature vector with the lab-type flag and passes the joint representation through a *tanh*-activated hidden dense layer (*units* = *n_feat*). A subsequent *soft-max* layer yields an attention vector $\boldsymbol{\alpha}$ of length *n_feat*; element-wise multiplication ($feat \times \alpha$) produces a context-weighted feature set that is forwarded to the classifier and retained for interpretability through trained attention extractor (*att_extractor*) enabling context-specific feature-importance analysis for pre- and post-G-CSF samples.

The downstream classifier comprises four fully connected layers with *leaky-ReLU* or *ReLU* activations, L2 weight regularization ($\lambda = 0.02$) on the first two layers, and batch normalization follows the initial block. Dropout rates, 0.1–0.2 are used sequentially to mitigate overfitting. Finally, a *sigmoid node* outputs the mobilization probability.

In a second model trained on CIBMTR donor dataset (n = 19,207), the architecture was deepened with six layers with decreasing units, each using leaky ReLU activations and followed by dropout (0.1–0.2), L2 regularization ($\lambda = 4.5 \times 10^{-6}$) and batch normalization where appropriate. This hierarchical structure allows the network to capture non-linear relationships among features while maintaining robustness against overfitting.

Both networks were trained using the *Adam optimizer* (*learning_rate* = $1 \times 10^{-4}$) with a binary-cross-entropy loss. Label smoothening was incorporated to reduce overconfidence in predictions. Model training was performed on the balanced training set with early stopping on the validation performance to minimize overfitting. Performance was monitored using ROC-AUC and binary accuracy metrics.

**Model evaluation and explainability**

For training and testing TabPFN, we used an 80:20 stratified train-test split to maintain class proportions. Model performance was then evaluated on a hold-out test set comprising 20% of the data. Importantly, because TabPFN necessitates minimal hyperparameter optimization, we were able to reserve the entire 20% hold-out set for independent testing. For the attention-aware model, we used an 80:20 train-test split; out of the 20% we carved 80% for validation and another 20% were utilized for testing. Specifically, the dataset was partitioned with two successive stratified random splits (random_state = 42), yielding 80% for model training, 16% for validation, and an untouched 4% hold-out set for final testing; all preprocessing/resampling was fitted on the training set only, with hyperparameters chosen on the validation set and the test set used once for final evaluation.

We assessed several performance metrics on the test set, including the area under the receiver operating characteristic curve (ROC–AUC), overall accuracy, precision, recall, F1-score and Mathews Correlation Coefficient (MCC), defined as:

$$MCC = \frac{TP \times TN - FP \times FN}{\sqrt{(TP + FP)(TP + FN)(TN + FP)(TN + FN)}} \quad (1)$$

Here, TP, FP, TN, and FN denote true positives, false positives, true negatives, and false negatives, respectively. The MCC is a correlation coefficient ranging from −1 (complete disagreement) through 0 (chance-level) to +1 (perfect prediction).

To interpret how model predictions were made, we employed SHAP (SHapley Additive Explanations) analysis. SHAP assigns each feature an importance value for individual predictions, enabling a game-theoretic understanding of the contribution of each predictor to the model's output (4). The SHAP package (4) was used to compute SHAP values for TabPFN-based models to identify which donor features most strongly influenced the mobilization outcome predictions. These interpretations were based on top 200 donor instances for pre-G-CSF (baseline) data trained model, and all test set instances (n=119) for the post-mobilization trained TabPFN model.

**Evaluation of attention mechanism performance**

To assess the quality and context sensitivity of the attention mechanism, we employed two complementary metrics: attention entropy and domain-specific attention divergence. Attention entropy, computed for each sample as

$$H(\alpha) = -\sum_{i=1}^{n} \alpha_i \log(\alpha_i) \tag{2}$$

where $\alpha = [\alpha_1, \alpha_2, \alpha_3, \dots, \alpha_n]$ denotes the attention weights assigned to the $n$ input features for a given sample. This quantifies the sharpness or focus of the attention distribution over input features. Lower entropy values indicate concentrated attention on a smaller subset of predictors, suggesting higher model interpretability and stronger feature selectivity. In contrast, higher entropy implies diffuse attention across many features, reflecting model uncertainty or weak discriminative preference. Moderate entropy values, as observed in our study, suggest a balance in which the model selectively attends to informative features while maintaining flexibility, supporting both interpretability and generalizability.

To assess domain adaptability, we computed the Jensen–Shannon (JS) divergence between the mean attention vectors for pre- and post-G-CSF samples. A higher JS divergence reflects more distinct attention patterns across lab types, confirming the network's ability to modulate feature importance based on sampling context. Feature-wise differences in attention were also analyzed to identify predictors differentially emphasized between domains.

**Supplementary Table 1:**

| | **CBC and Demographic Features** |
|---|---|
| 1. | Gender |
| 2. | Age |
| 3. | White Blood Cells (WBC) |
| 4. | Hemoglobin (Hgb) |
| 5. | Hematocrit (Hct) |
| 6. | Platelet Count |
| 7. | Neutrophil Percentage |
| 8. | Lymphocyte Percentage |
| 9. | Monocyte Percentage |
| 10. | Eosinophil Percentage |
| 11. | Neutrophil Absolute |
| 12. | Lymphocyte Absolute |
| 13. | Monocyte Absolute |
| 14. | Eosinophil Absolute |

**Supplementary Figure 1:**

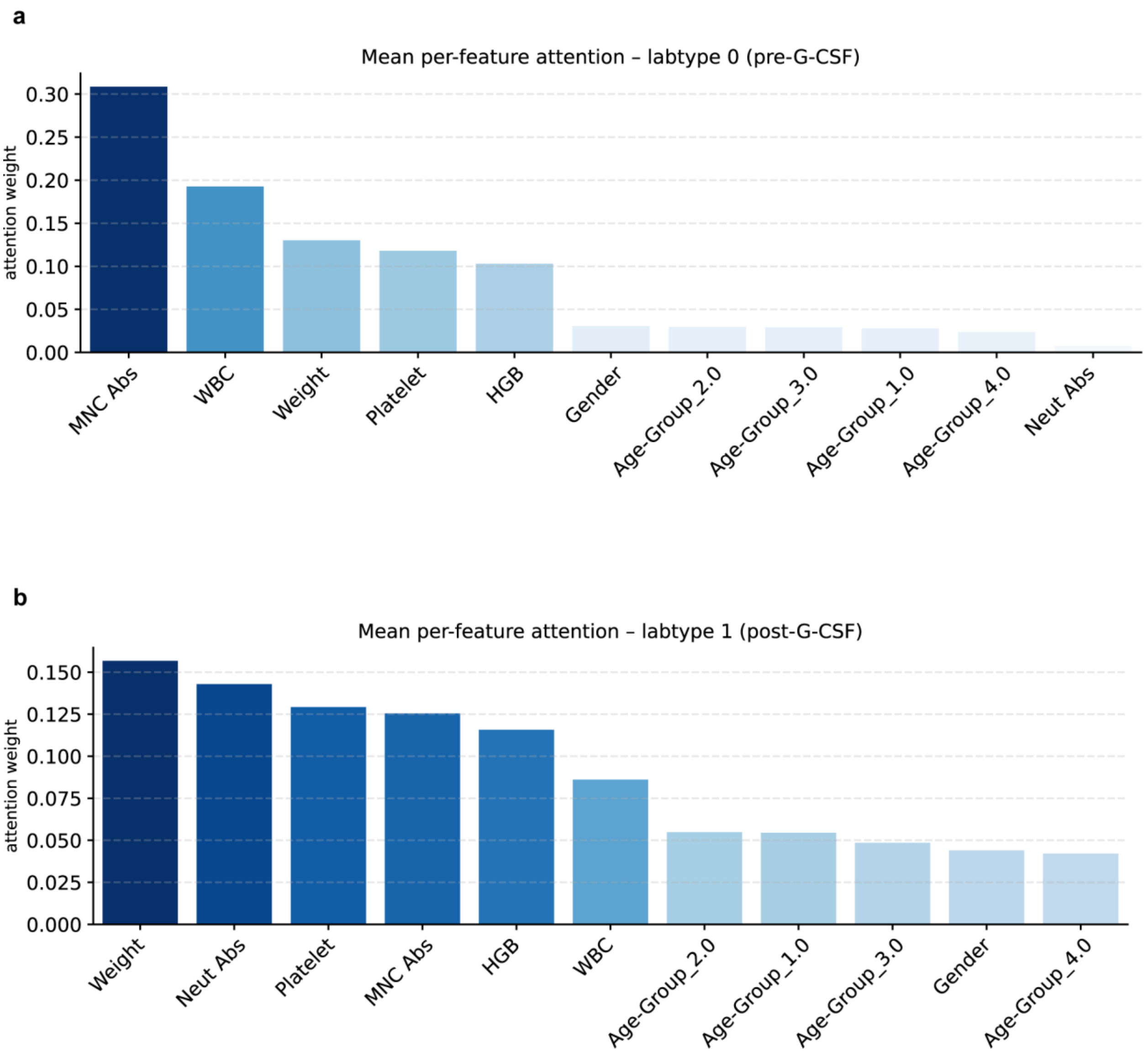


**Figure 1. Feature-wise attention profiles across lab contexts. (a)** Mean per-feature attention weights for labtype 0 (pre-G-CSF), showing dominant contributions from absolute mononuclear cell count (MNC Abs), donor weight, platelet count, and hemoglobin (HGB). **(b)** Mean per-feature attention weights for labtype 1 (post-G-CSF), with donor weight, platelet count, and MNC Abs receiving the highest attention.

**Supplementary Figure 2:**

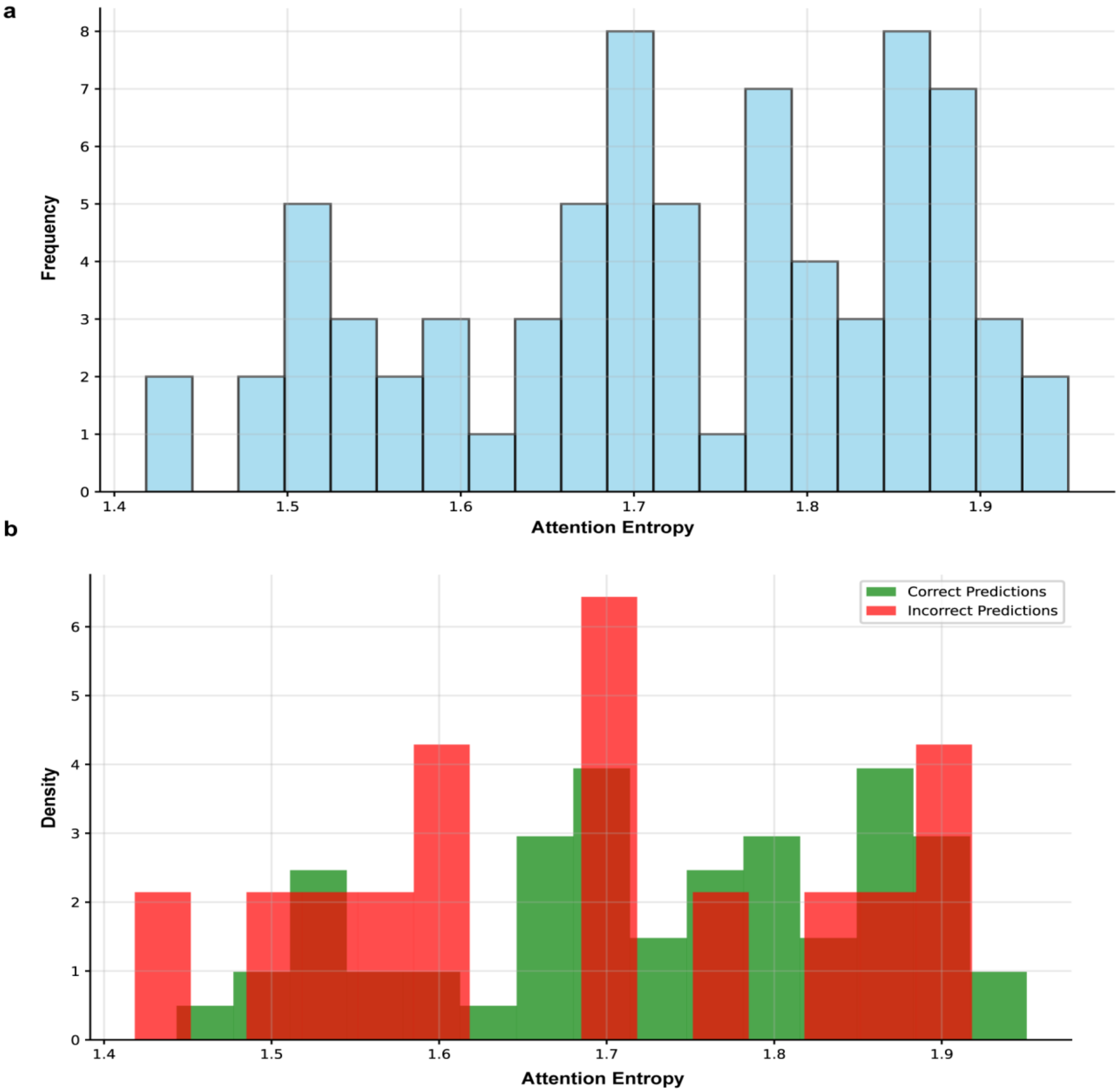


**Figure 2. Distributions of attention entropy across prediction outcomes. (a)** Histogram showing the overall distribution of attention entropy values across all samples. The mean attention entropy was 1.73 (natural log scale), suggesting that the model neither distributed its attention uniformly across all features nor overly concentrated it on a single input. **(b)** Density histogram stratified by prediction accuracy, highlighting the attention entropy distribution for correctly classified (green) versus incorrectly classified (red) samples. Entropy distributions were comparable between correct and incorrect predictions (1.73 vs. 1.69), indicating that selective attention was a general property of the model rather than specific to only high-confidence cases. Furthermore, negligible correlation between attention entropy and prediction confidence (Pearson's r=0.019, p=0.87) implies that focused attention was not merely a proxy for model certainty, but an independent learned mechanism. Jensen–Shannon divergence between attention distributions in pre- and post-G-CSF samples was 0.274.
